# Single-cell transcriptomic assessment of cellular phenotype stability in human precision-cut lung slices

**DOI:** 10.1101/2021.08.19.457016

**Authors:** Nichelle I. Winters, Chase J. Taylor, Christopher S. Jetter, Jane E. Camarata, Austin J. Gutierrez, Linh T. Bui, Jason J. Gokey, Matthew Bacchetta, Nicholas E. Banovich, Jennifer M.S. Sucre, Jonathan A. Kropski

**Affiliations:** Division of Allergy, Pulmonary and Critical Care Medicine, Department of Medicine, Vanderbilt University Medical Center, Nashville, TN; Division of Neonatology, Department of Pediatrics, Vanderbilt University Medical Center, Nashville, TN; Translational Genomics Research Institute, Phoenix, AZ; Department of Thoracic Surgery, Vanderbilt University Medical Center, Nashville, TN; Department of Cell and Developmental Biology, Vanderbilt University, Nashville, TN; Department of Veterans Affairs Medical Center, Nashville, TN

## Abstract

Precision-cut lung slices (PCLS) are increasingly utilized for *ex vivo* disease modeling, but a high-resolution characterization of cellular phenotype stability in PCLS has not been reported. Comparing the single-cell transcriptomic profile of human PCLS after five days of culture to freshly isolated human lung tissue, we found striking changes in endothelial cell and alveolar epithelial cell programs, reflecting both injury and pathways activated in static culture, while immune cell frequencies and programs remained largely intact and similar to the native lung. These cellular dynamics should be considered when utilizing PCLS as a model of the human lung.

## INTRODUCTION

Diseases of the respiratory system are a leading cause of death worldwide [1]. *Ex vivo* and high fidelity models of human pulmonary biology are essential for the development and clinical translation of novel, disease-modifying therapies for lung diseases. Precision-cut lung slices (PCLS) have emerged as a promising disease modeling platform as in theory, the cellular diversity, 3-dimensional architecture, and cell-cell/cell-matrix interactions are maintained. PCLS are amenable to long term culture (up to 15 days), tolerate cryopreservation, and have been generated from both diseased (COPD, asthma, interstitial lung disease) and healthy human lungs [2]. While cell phenotypes have been characterized using standard histological approaches [3], there have been no high-resolution studies of how cellular transcription programs evolve during PCLS culture. Understanding the molecular changes that occur in PCLS culture is critical to inform which applications and questions PCLS can be effectively utilized to study.

## METHODS

Detailed methods provided in online supplement.

## RESULTS

We generated PCLS from the right middle lobe of a human donor without pre-existing lung disease and cultured the slices for 5 days. After 5 days of culture, the lung slices appeared histologically similar to time 0 (Figure 1, A-C). We pooled 12 PCLS after 5 days of culture to generate a single-cell suspension, performed single cell RNA sequencing (scRNA-seq), and compared the transcriptomic profile from the PCLS to a pooled population of freshly isolated single cells from 11 human donor lungs (supplemental tables). All cell clusters derived from fresh tissue were also identified in 5 day-PCLS with the exception of two secretory cell subsets (Figure 1D, F). Expression of canonical markers in parenchymal cells, while detectable, was lower in PCLS compared to fresh tissue. This was most notable in the endothelial (*ACKR1, CA4, HEY1*), AT1 (*RTKN2, AGER, HOPX*) and AT2 (*LAMP3, ABCA3, SFTPC*) populations (Figure 1, E).

**Figure 1.**
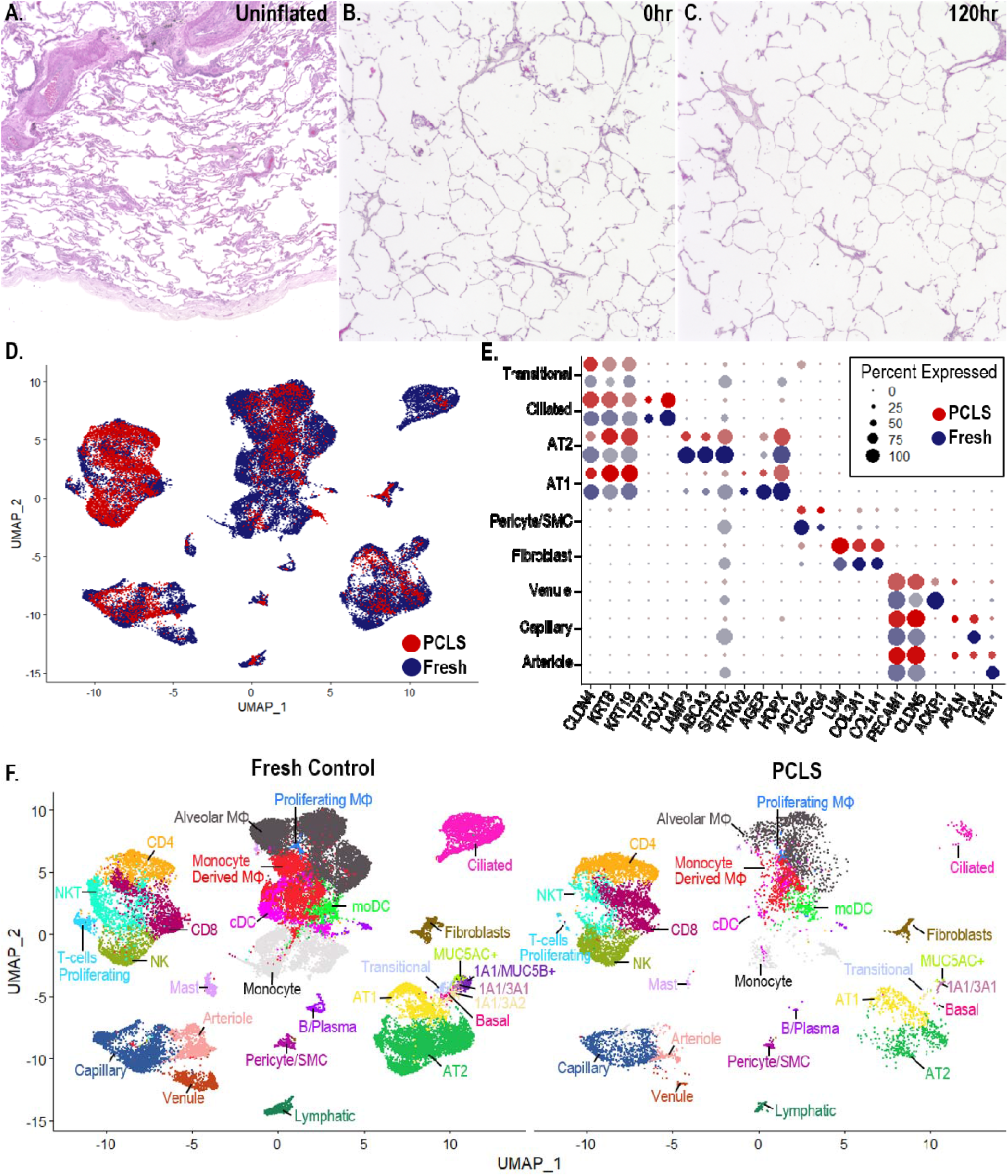
Cellular diversity in PCLS is maintained after 5 days of culture. A) H&E stain of adjacent tissue segment of un-inflated donor lung used to generate PCLS. B) H&E stain of tissue block after inflation with agarose prior to culture. C) H&E stain of PCLS after 120hrs of culture appears similar to time 0. D) Integrated UMAP demonstrates similar clustering between Fresh (blue) and PCLS (red) cells. E) Dot plot comparing canonical marker expression of non-immune cells. Color intensity indicates average expression level. F) Annotated UMAP of Fresh Control versus PCLS.

The process of generating PCLS inherently disrupts the alveolar-capillary network in addition to macrovascular endothelial structures. We observed that PCLS endothelial cells upregulated a subset of tip cell markers including APLN, CD34, DLL4, KDR, PLK2, PLAUR, LCP2, ROBO4, CXCR4, LXN, while others were found at lower levels in PCLS culture including FLT4 and UNC5B [4,5]. In contrast, markers of stalk cells (ACKR1, SELP, VWF, TEK), another important cell type for functional branching angiogenesis, were found at lower levels in PCLS endothelial cells suggesting induction of genuine angiogenesis may not account for these changes in gene expression [6]. Endothelial cell gene expression is known to be modulated by flow conditions. Laminar flow, shear stress and static culture have all been linked to activation or downregulation of specific gene groups which often overlap with angiogenesis markers [7,8]. Indeed, in the PCLS, a static culture state, genes associated with flow (laminar or shear) were downregulated (THBD, TFPI, KLF2, ID, VWF, TEK) and genes previously linked to static culture conditions were upregulated (CXCR4, LXN, EFNA1) compared to freshly isolated control lung tissue (Figure 2A, supplemental tables 1-3).

**Figure 2.**
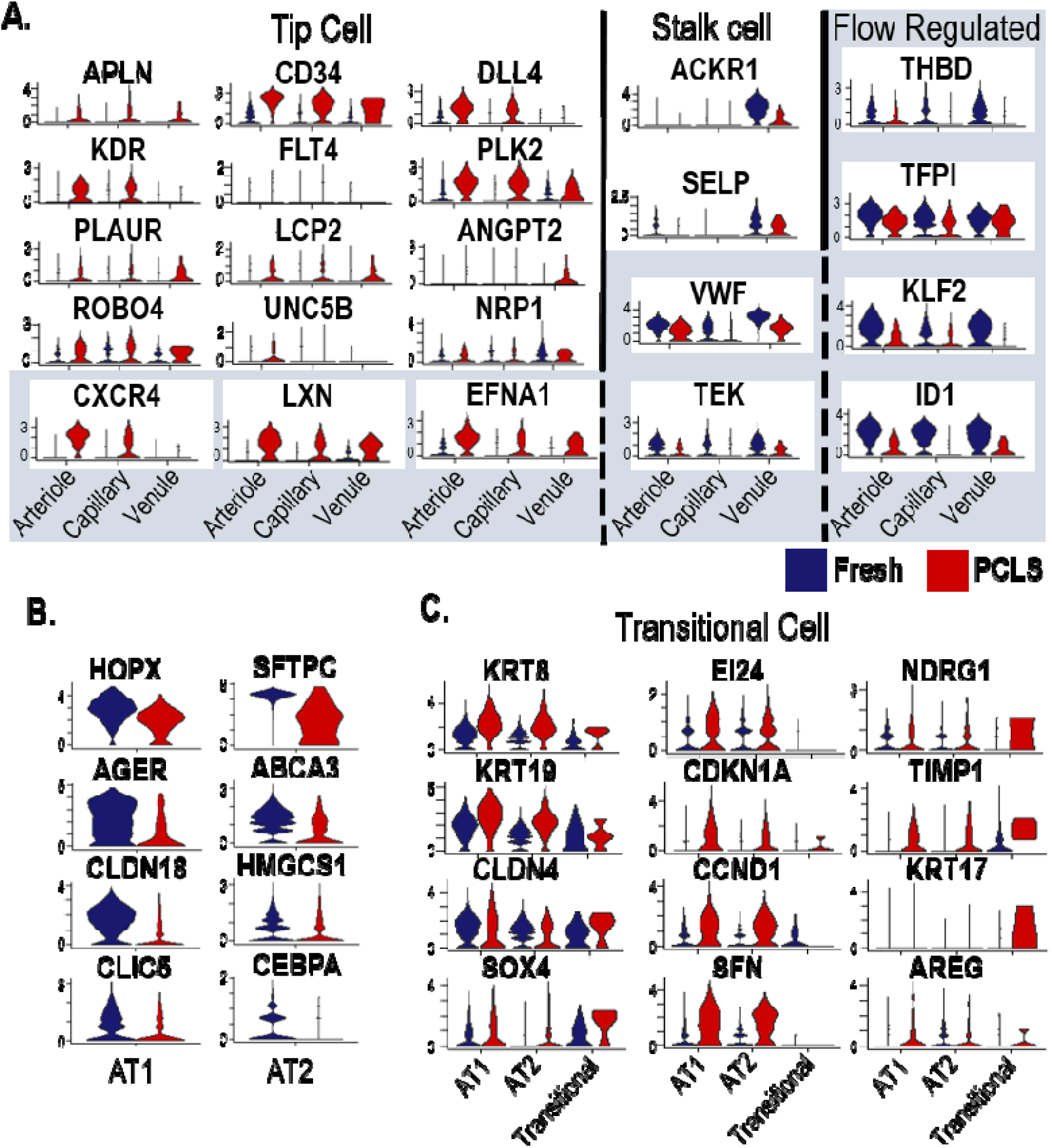
Gene expression profile changes in endothelial and alveolar epithelial cells between PCLS and Fresh cells. A) Violin plots of arteriole, capillary and venule endothelial cell clusters. Flow regulated gene expression is denoted with a gray background. A subset of flow regulated genes overlap with tip cell and stalk cell markers (columns). Tip cell markers are increased by PCLS culture but there is not a concurrent upregulation of stalk cell markers. Markers of static culture are upregulated and shear stress markers are downregulated in PCLS endothelial cells. B) Violin plots of AT1 and AT2 marker genes demonstrate downregulation after PCLS culture. C) Transitional cell marker genes are upregulated in PCLS culture in AT1 and AT2 cells.

Similar to endothelial cells, alveolar epithelial cells (AEC) are constantly exposed to dynamic mechanical forces during respiration which modulates cell differentiation and behavior. While PCLS and Fresh Control AT1 and AT2 cells cluster together, there is a downregulation of canonical markers of terminal AEC states among PCLS cells (Figure 2B) compared to freshly isolated AECs. Concurrently, there is an upregulation of markers of transitional cells in PCLS culture (Figure 2C, supplemental tables 4-6). Transitional cells are an intermediate cell type between AT2 and AT1 cells and are induced by multiple stimuli including injury and stretch which are both features of PCLS culture [9,10].

A significant limitation of many organoid/*ex-vivo* lung disease models is the inability to study interactions involving innate and adaptive immune cells due to poor survival in cell-culture conditions. In contrast, a substantial proportion of the cells recovered from the PCLS were viable immune cells (Figure 1D, F) and could be identified by their expression of canonical markers (Figure 3A). While all major immune cell types were recovered from PCLS, their relative frequencies differed compared to fresh tissue. This was most evident in alveolar macrophages which were proportionately reduced while NK cells comprised a larger proportion of recovered immune cells in PCLS (Figure 3B). This loss of alveolar macrophages over time was also observed by immunoflourescent analysis and is at least partially due to cell death as evidenced by TUNEL staining (Figure 3 C-D). Despite the alterations in relative population numbers, the changes in the gene expression profiles of alveolar macrophages and NK cells were predominantly shared across all myeloid and lymphoid cell types, respectively. In contrast, monocytes and NKT cells had a large number of uniquely differentially expressed genes suggesting a cell type specific alteration occurs in PCLS culture of these cell types (Figure 3 E-F, Supplemental Tables 7-17, Supplemental Figure 1).

**Figure 3.**
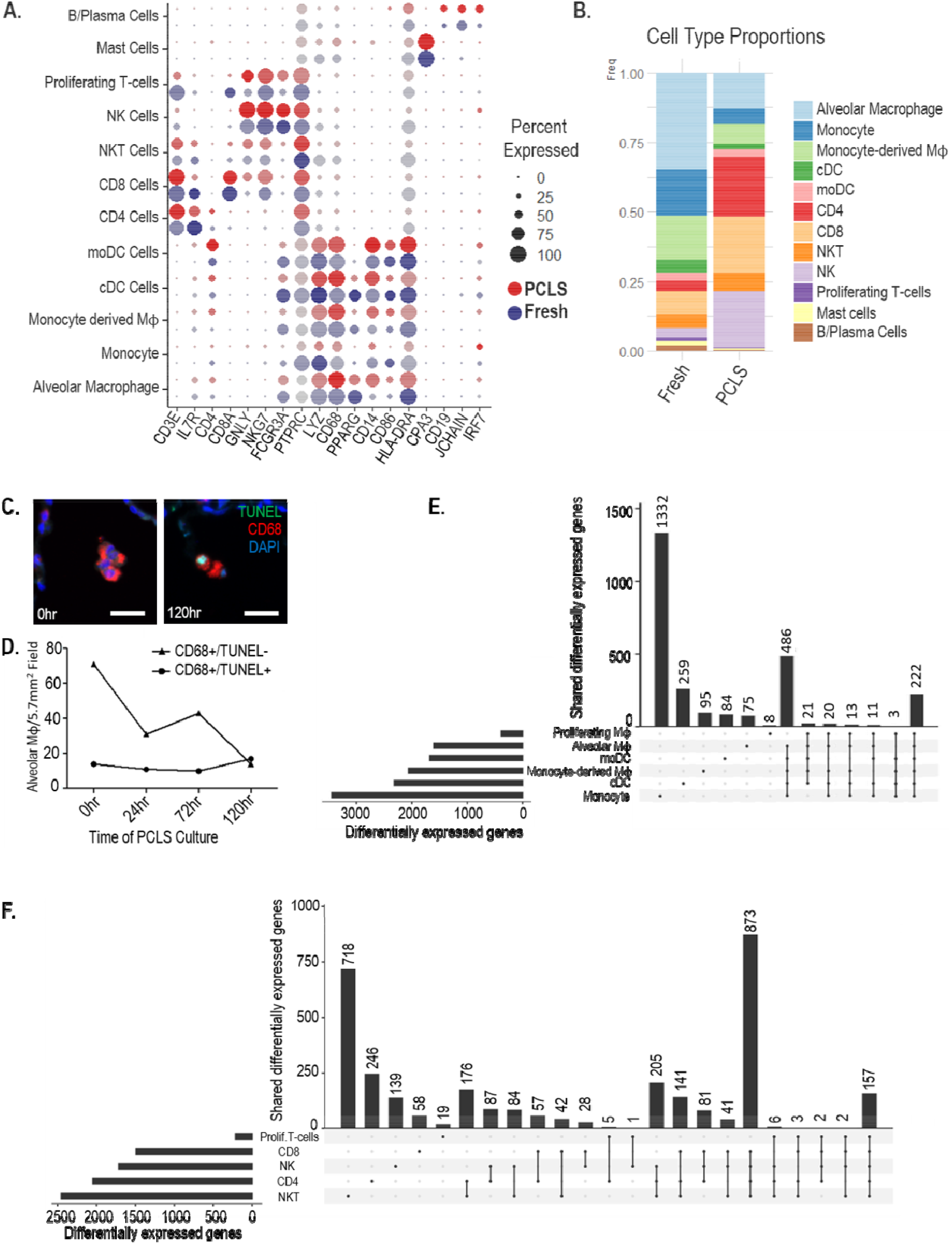
Immune cell transcriptional changes induced by PCLS culture. A) DotPlot demonstrating major immune cell class markers are maintained in PCLS culture. B) Immune cell proportions recovered from Fresh and PCLS tissue. C-D) CD68 (red) positive alveolar macrophages at time 0 are more abundant and predominantly TUNEL (green) negative. After 120hrs of culture, TUNEL positive alveolar macrophages are proportionately increased and overall numbers of alveolar macrophages decreased. E) Upset plot of shared differentially expressed genes between fresh and PCLS cells of the myeloid lineage. F) Upset plot of shared differentially expressed genes between fresh and PCLS cells of the lymphoid lineage. Significance level for differentially expressed genes is p-value<0.05.

## DISCUSSION

These data are the first report of successful scRNA-seq on PCLS, demonstrating the feasibility of this technique as well as providing a first look at broad changes that occur during PCLS culture. Overall, there is a remarkable conservation of pulmonary cellular phenotypes in PCLS culture and this technique is appropriately heralded for its fidelity to the *in vivo* condition in comparison to other *in vitro* techniques that incorporate only a subset of cell types. However, there are important changes in key cell types. In particular, with long term culture, PCLS induces changes in endothelial cell phenotypes which resemble static culture states, while AT1 and AT2 cells appear to undergo a partial de-differentiation and adopt a transitional cell-like phenotype - these substantial phenotypic changes should be anticipated when utilizing PCLS for long term studies of endothelial and epithelial function. Finally, PCLS culture conditions induce a broad but shared transcriptomic shift across the majority of immune cell types though there are cell type specific changes both in cellular proportions as well as gene expression profiles that especially affect monocytes, alveolar macrophages, NK and NKT cells. By taking these changes into account, effective experiments can be designed that utilize this powerful model system of the human lung.

## ACKNOWLEDGEMENTS

This study was supported by the National Institutes of Health R01HL145372(JAK/NEB), the Doris Duke Charitable Foundation (JAK), T32HL094296 (NIW), K08HL143051(JMSS), R01HL140231(MJB).

