## Supplement for "Single-cell transcriptomic assessment of cellular phenotype stability in human precision-cut lung slices"

**Supplemental Methods**

Subjects and samples: Lung tissue was isolated from lungs declined for transplantation at the time of organ donation. This study was approved by the Vanderbilt University Institutional Review Board (IRB #171657, 192004, 060165).

Generation and culture of PCLS: Right middle lobe bronchus was identified, cannulated and injected with ~35cc of 2% low melting point agarose in DMEM/F12 (Gibco 11320033) with 1X Anti-Anti (Gibco 15240096) at 37°C. After inflation, the bronchus was clamped off and the entire lobe was cooled at 4°C for 1 hr. A 1cm/1cm cube was excised from the inflated segment and 500μM PCLS were generated with a vibratome (Leica). PCLS were washed with sterile PBS and then washed 5X 30 min in sterile DMEM/F12 (Gibco). PCLS were then incubated in DMEM/F12 + Anti-Anti for 120 hours. Sections were isolated at 24 hour timepoints, fixed in formalin and embedded in paraffin for sectioning and histological staining.

Immunofluorescent staining and quantification of CD68+ cells: 5μm sections of paraffin embedded PCLS at time 0, 72 and 120 hrs of culture were deparaffinized and rehydrated. Sodium citrate antigen retrieval was performed followed by blocking in 10% donkey serum. Sections were labeled with anti-CD68 (SCBT sc-20060, 1:100) followed by donkey anti-mouse 647 secondary (Invitrogen A-31571), TUNEL assay (Roche, 11684795910), and DAPI. The entire section was captured at 10x magnification using automated tiling on a Keyence BZ-X800. The center of the section was then identified and marked and a 5X5 grid at 20X magnification was captured around the center of the PCLS. All CD68+ cells within the alveolus within this 5X5 grid (5.7mm^2^) were counted and noted for presence or absence of nuclear TUNEL positivity.

scRNA-seq library preparation and sequencing: Single-cell suspensions were generated by digesting 2 grams of fresh tissue or pooling 12 PCLS and performing an enzymatic digest using a combination of collagenase I and dispase (1 mg/ml each) as previously described (*1*). Agarose appears to be visually filtered out with the 100μM filter step. Library preparation was performed using the 10X Chromium 5’ v1 kit targeting 20,000 cells. Library sequencing was performed on an Illumina Novaseq6000 targeting 50,000 reads/cell as previously described (*1*). Demultiplexing was performed using CellRanger v5.0. Freshly processed scRNA-seq library from this same donor, as well as raw sequencing data from our previously published dataset (GSE135893) were demultiplexed and aligned using CellRanger v5.0.

scRNA-seq analysis. After alignment and demultiplexing, fresh and PCLS-cultured libraries were merged, underwent quality control filtering (excluding cells with <750 genes, <0.1% or >15% mitochondrial reads), followed by batch-correction (fresh samples vs. PCLS) using the SCTransform-integration workflow in Seurat v4 (*2*), followed by joint clustering, uniform-manifold approximation and projection (UMAP) (*3*), and cell-type annotation using canonical markers informed by our previous publication (*1*) and other recently reported lung atlases (*4*–*9*). Cell-type specific differential expression was performed using the negative binomial test using a logFC threshold of 0.25 and adjusted p<0.05 as a significance threshold. All code used for integration and analysis of this dataset is available at <https://github.com/KropskiLab/PCLS>.

Data availability: Raw genomic data are available at the Gene Expression Omnibus (GEO) GSE181388 (PCLS and matching fresh sample) and GSE135893 (fresh control samples).

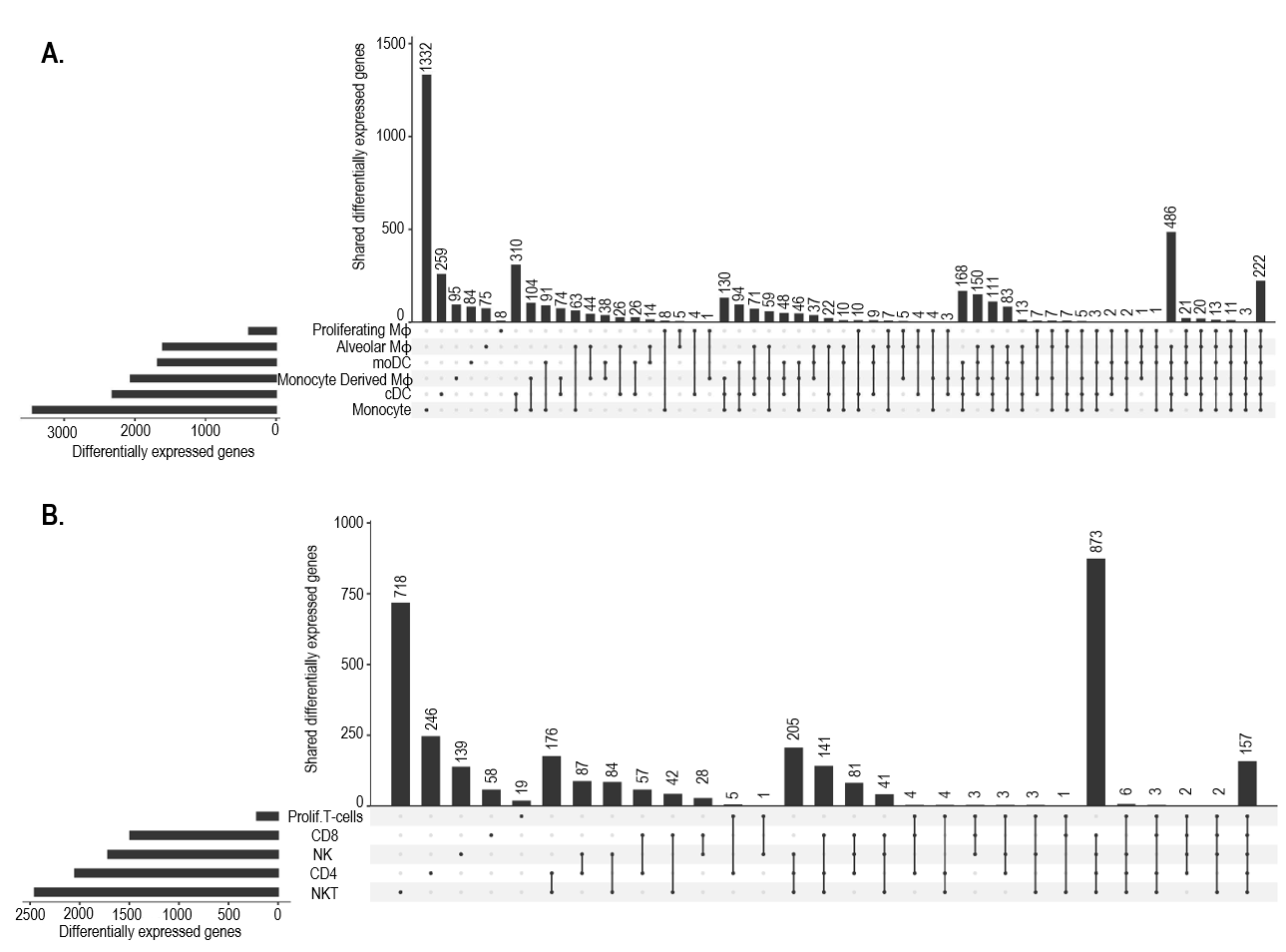

**Supplemental Figure 1: Complete Myeloid and Lymphoid Upset Plots.** A) Upset plot of shared differentially expressed genes among the myeloid lineage. B) Upset plot of shared differentially expressed genes among the lymphoid lineage. Significance level for differentially expressed genes is p-value<0.05.
